# The Kinetic Landscape of Nucleosome Assembly: A Coarse-Grained Molecular Dynamics Study

**DOI:** 10.1101/2021.02.14.431121

**Authors:** Giovanni B. Brandani, Cheng Tan, Shoji Takada

## Abstract

The organization of nucleosomes along the Eukaryotic genome is maintained over time despite disruptive events such as replication. During this complex process, histones and DNA can form a variety of non-canonical nucleosome conformations, but their precise molecular details and roles during nucleosome assembly remain unclear. In this study, employing coarse-grained molecular dynamics simulations and Markov state modeling, we characterized the complete kinetics of nucleosome assembly. On the nucleosome-positioning 601 DNA sequence, we observe a rich transition network among various canonical and non-canonical tetrasome, hexasome, and nucleosome conformations. A low salt environment makes nucleosomes stable, but the kinetic landscape becomes more rugged, so that the system is more likely to be trapped in off-pathway partially assembled intermediates. We find that the co-operativity between DNA bending and histone association enables positioning sequence motifs to direct the assembly process, with potential implications for the dynamic organization of nucleosomes on real genomic sequences.

## Introduction

Nucleosomes are the structural units of chromatin, each consisting of ∼147 base pairs of DNA wrapped around a protein histone octamer made of H3/H4 tetramer and two H2A/H2B dimers [1,2]. Nucleosomes not only enable the efficient compaction of the long Eukaryotic genomic DNA into the nucleus, but also influence key biological process by controlling the access to DNA by other proteins [3], for example during transcription [4]. Such important functions rely on the establishment of precise epigenetic patterns of nucleosome positions, histone variants and histone modifications, which mark the distinction between active and repressed genes, and between different genomic regions, e.g. promoters and gene bodies [5]. In order to preserve the identity of each cell, essential in multi-cellular organisms, this genomic organization has to be maintained [6] despite the continuous disassembly of nucleosomes over the course of DNA transcription [7] and replication [8]. However, the molecular details of nucleosome assembly and disassembly have not been yet fully elucidated.

*In vitro* experiments showed that the salt-induced disassembly of nucleosomes starts with DNA unwrapping and it is then followed by the progressive loss of the two H2A/H2B dimers [9], therefore forming firstly an asymmetric hexasome, and finally a tetrasome. Assembly proceeds in the opposite direction, starting with the binding of a H3/H4 tetramer, then followed by the binding of a H2A/H2B dimer on each side [10]. This order of events is due to the specific geometry of the nucleosome (with the H2A/H2B dimers surrounding the central H3/H4 tetramer), and to the more extensive network of protein-DNA hydrogen bonds on H3/H4 relative to H2A/H2B [1,11]. A similar assembly pathway was suggested to occur also *in vivo* [12], except that in this case the process is facilitated by histone chaperones limiting non-nucleosomal histone-DNA interactions [12], or by chromatin remodelers [13]. However, many experiments paint a picture of assembly (or disassembly) more complicated than a simple 2-step process involving tetrasomes, hexasomes and complete nucleosomes, highlighting the existence of various partially-assembled or non-canonical nucleosome structures, such as pre-nucleosomes [14], right-handed nucleosomes [15], remodeler-associated fragile nucleosomes [16], and partially opened nucleosomes [17]. Sub-nucleosomal conformations are also widely distributed across the genome *in vivo* [18], and their formation has been shown to depend in part on the underlying DNA sequence [19]. How such structures form and interconvert between each other remains unclear.

Characterizing the precise molecular details of nucleosome intermediates and the factors controlling their formation, e.g., salt concentration and DNA sequence, would greatly aid our understanding of how chromatin organization is established and maintained. In order to address these questions, we ran extensive coarse-grained molecular dynamics (MD) simulations of nucleosome assembly under different conditions. We consider a simple system made of the strong nucleosome-positioning 601 DNA sequence [20], a H3/H4 tetramer and two H2A/H2B dimers. By analyzing our MD trajectories with Markov state modeling [21], we reveal a complex kinetic landscape characterized by many metastable structures, some of which correspond to previously proposed non-canonical nucleosomes. Our simulations show that A/T nucleosome positioning signals on the 601 DNA sequence direct the binding of the H2A/H2B dimer on the optimal DNA region, so that the dimer is then poised to easily bind the H3/H4 tetramer interface. This effect, which is observed only on the A/T-rich side of the 601 DNA, reveals a highly sequence-dependent assembly process, suggesting a potential mechanism allowing genomic sequences to directly control nucleosome organization *in vivo* [22]. We also investigated the influence of salt on the kinetics, finding that successful nucleosome assembly from a tetrasome is faster at low salt concentration, but that the system is also more likely to remain stuck in off-pathway kinetic traps. This explains the necessity to assemble nucleosomes via salt dialysis [23], or via the addition of chromatin remodelers [14]. The assembly/disassembly mechanism itself is also a function of salt concentration: while at low or intermediate salt the H2A/H2B dimers bind/unbind to/from the H3/H4 tetramer co-operatively with DNA wrapping, unbinding occurs only after DNA unwrapping at high salt. Overall, our results establish clear mechanistic relations between environmental conditions, DNA sequence, and the dynamics of nucleosome assembly.

## Results

### Modeling nucleosome assembly dynamics

In our MD simulations, the DNA is modeled using the 3SPN2.C coarse grained model [24], where each nucleotide is represented by three beads centered on the base, sugar and phosphate groups. The histones are modeled according to the AICG2+ structure-based model [25], where each residue is represented by one bead centered on the Cα atom. This and similar coarse-grained models have been successfully applied to study complex processes such as nucleosome sliding [26–29], unwrapping [30–32], and assembly [33], at a fraction of the computational cost required for all-atom simulations [34–37]. Notably, the 3SPN2.C model can capture the sequence-dependent elasticity of DNA [24], and it has been shown to correctly predict the affinity of nucleosome formation for different sequences [32], making this model suitable to investigate the effect of sequence on the assembly kinetics. Further details on the coarse-grained model and its parametrization are provided in the Methods section.

Our simulation system consists of the 147-bp nucleosome-positioning 601 DNA [20], one histone H3/H4 tetramer, and two copies of histone H2A/H2B dimers. The 601 sequence was chosen for its widespread use in experiments and for their characteristic positioning motifs asymmetrically distributed around the nucleosome, allowing us to study their role during assembly. Specifically, 10-bp periodic A/T base steps are more frequent on the left side of 601 (called the beta side in Ref. [17], see Fig 1A), leading to enhanced nucleosome unwrapping on the right side relative to the left (i.e. asymmetric unwrapping) [9,38].

**Fig 1.**
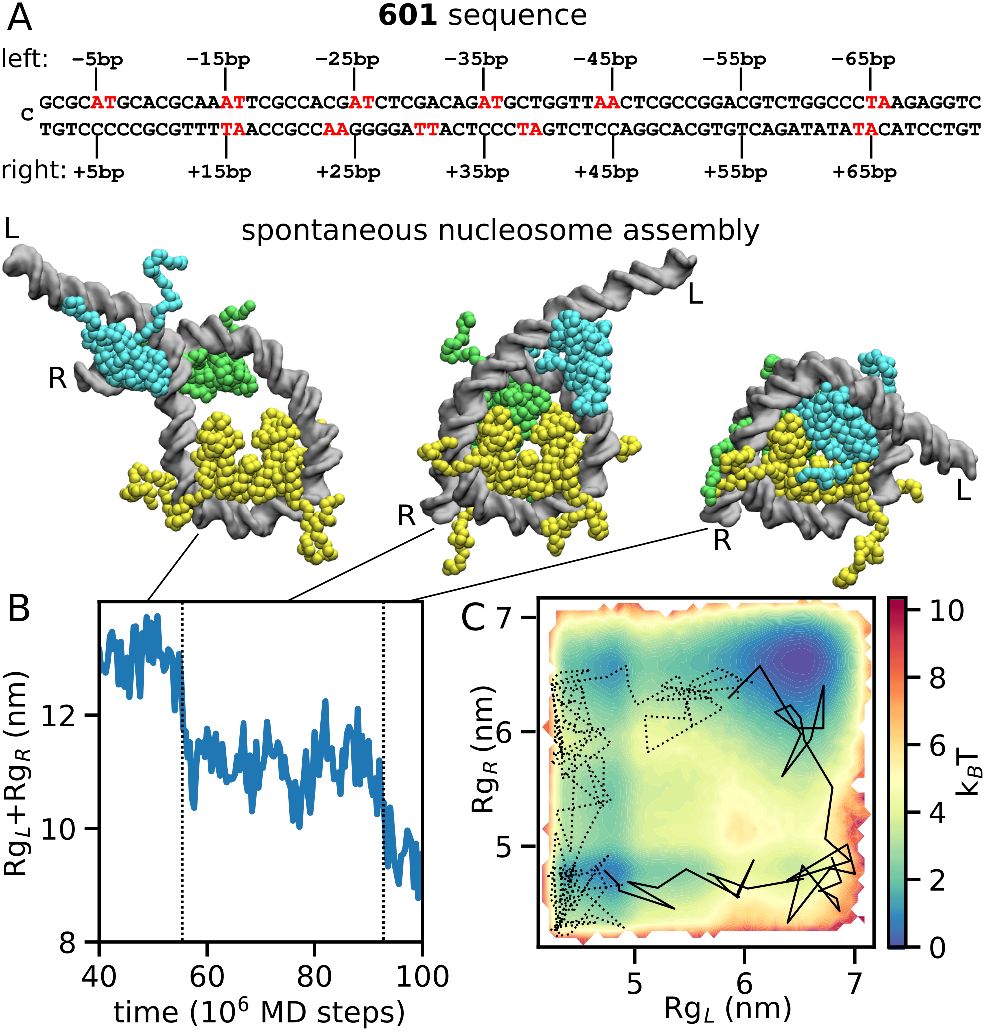
Nucleosome assembly simulation for the 147-bp 601-sequence at 400 mM. (A) The 601 DNA sequence. The left (0 to -73 bp) and right (0 to +73 bp) halves are shown on the first and second lines, relative to the dyad position at 0 bp (the left end), highlighting the locations of the A/T periodic motifs in red. (B) Timeline of the sum of radii of gyration of the left and right halves of DNA (Rg_L_+Rg_R_) during the formation of a complete nucleosome starting from a tetrasome. In the 3 snapshots corresponding to tetrasome (left), hexasome (center) and nucleosome (right), the DNA is shown in grey, the H3/H4 tetramer in yellow, the H2A/H2B dimer binding to the left side of 601 in cyan, and the H2A/H2B dimer binding to the right in green (throughout the paper, the same color code is used). The left and right DNA entry/exits are indicated by the letters L and R respectively. (C) The same trajectory as in panel B (at 400 mM, solid line) shown in the two dimensions, Rg_L_ and Rg_R_, together with one successful trajectory at 200 mM (dotted line). The free energy landscape at 400 mM calculated by the Markov state model is also plotted.

We performed nucleosome MD simulations for 10^8^ MD steps at 400 mM mono-valent ion concentration starting from the tetrasome conformation, in which the H3/H4 tetramer is bound on the 601 DNA at the optimal position and both of H2A/H2B dimers are unbound from the tetramer but bound on DNA at different positions depending on the simulation run (see the left snapshot in Fig 1B as an example). Out of 215 runs, we found only one trajectory that reached to the complete nucleosome, which is shown in Fig 1B and S1 Movie. As highlighted by the timeline of the sum of the radii of gyration of the left and right DNA sections (Rg_L_+Rg_R_), in this successful trajectory assembly occurs in two steps: firstly, the right half of DNA bends together with the binding of one H2A/H2B dimer to the tetramer (central cartoon in Fig 1B), then, the left half of DNA folds concomitantly with the assembly of the other H2A/H2B dimer to form the complete nucleosome (right cartoon). In Fig 1C, the same trajectory is projected on the 2-dimensional free energy landscape (obtained by the Markov state model as described below) along the left and right radii of gyration of DNA. The free energy landscape also shows that the left side of the 601 DNA bends more easily (i.e., with a lower free energy cost) than the right side; this asymmetry is discussed later in the results. Other trajectories stopped at various apparently metastable states, due to the limitation of the simulation time.

Starting from the same tetrasome conformations used at 400 mM, we also performed 215 simulations at 200 mM and 300 mM, finding respectively 20 and 7 cases of complete nucleosome assembly (a representative trajectory at 200 mM is shown in Fig 1C). We will discuss the salt concentration dependence in more details later in the results.

### Kinetic landscape of nucleosome assembly via Markov state modeling

In order to characterize the complete landscape and kinetics of nucleosome assembly and disassembly, we generated a Markov state model (MSM) of nucleosome dynamics using 1000 independent simulation runs of 10^8^ MD steps each at 400 mM salt, starting from a diverse set of partially or fully assembled nucleosomes (215 of these are the same described in the previous section, see Methods for more details). The salt concentration of 400 mM was chosen so that neither the fully disassembled nor the fully assembled nucleosome conformations are too much preferred over the others, facilitating the characterization of the dynamics. To build an MSM, we first divide the conformational space into a set of *M* discrete microstates, each corresponding to a group of similar conformations, and then we learn from the unbiased MD trajectories the *M* x *M* transition probabilities to go from each state to another after a certain time interval, the so-called lag-time. Observing either complete assembly or disassembly events in a single trajectory is extremely rare, but the Markov state modeling approach allows us to combine many trajectories together to fully characterize the kinetics of the system, identifying the main long-lived conformations and the dominant pathways of nucleosome assembly. We generated an MSM with *M*=400 microstates and a lag-time of 4×10^6^ MD steps (see Methods section for more details, and S1 Fig for the convergence of the MSM timescales). To aid the interpretation of the model, we then further grouped the 400 microstates into 11 metastable basins associated to the 10 slowest relaxations of the MSM (obtained via PCCA clustering [39]). We obtained the transition probabilities between the basins from a coarse-grained hidden Markov state model derived from the full MSM [40].

Fig 2 summarizes the resulting kinetic landscape of nucleosome assembly and disassembly at 400 mM, displaying the identified 11 metastable basins and transition probabilities between these basins. Of the 11 basins, we find one state corresponding to the fully assembled canonical nucleosome (N), seven hexasome states (H_X_) and three tetrasome states (T, T_L_ and T_R_). In S2 Fig we show that the TICA coordinates [41] used to generate the MSM can distinguish well these states, as a further indication of their metastability. The three tetrasome states are distinguished by the positions of the unbound H2A/H2B dimers. In the T state one dimer is on the left side of the unwrapped DNA while the other dimer is on the right side, remaining close to the target H3/H4 interface. On the other hand, in the T_L_ and T_R_ states both unbound dimers interact solely with either the left or right side of the unwrapped DNA, respectively. The seven hexasome structures are divided into two types, depending on whether one H2A/H2B dimer is bound to the left (H_L0_, H_L1_ and H_L2_) or the right (H_R0_, H_R1_, H_R2_ and H_R3_) binding interface of the H3/H4 tetramer (respectively wrapping the left or the right sides of the 601 DNA in the canonical nucleosome). Each of these two hexamer types can adopt different conformations depending on the location of the unbound H2A/H2B dimer. In the left hexasome H_L0_, the unbound dimer is either far away or it interacts only weakly with nucleosomal DNA. In H_L1_, the unbound dimer is located on the right side of the DNA relatively close to its optimal location in the complete nucleosome. In H_L2_, the unbound dimer bridges two DNA gyres while facing outside of the nucleosome, in a configuration that cannot easily lead to assembly. H_R0_, H_R1_ and H_R2_ are the respective right hexasome versions of those just described. Finally, in hexasome H_R3_ a H2A/H2B dimer binds to the right interface of the H3/H4 tetramer, but it is stabilized by the wrapping of the left side of the 601 sequence. This metastable configuration is not found on the opposite side (there is no H_L3_ basin), suggesting that it originates from the marked propensity of the left side of 601 to bend. Notably, in the H_R3_ hexasome the DNA follows a right-handed super-helix, contrary to the canonical form that follows a left-handed one. For each of the 11 identified metastable states, we deposited as supplementary material 4 representative coarse-grained nucleosome configurations in PDB format (S1 Data). Researchers interested in investigating sub-nucleosome conformations in all-atom MD may use these as starting configurations following a previously-developed back mapping procedure [42].

**Fig 2.**
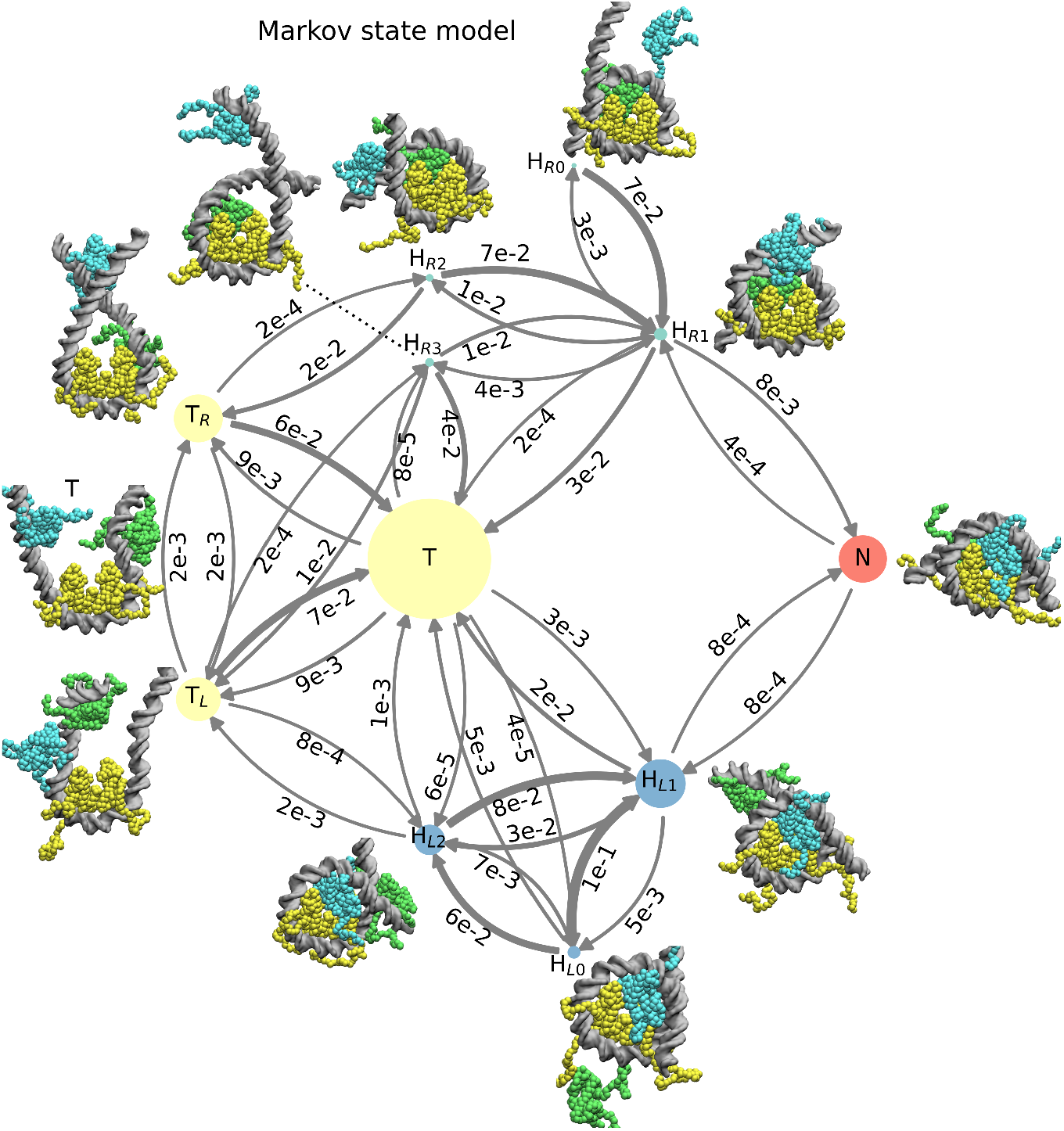
Kinetic landscape of nucleosome assembly and disassembly at 400 mM from the Markov state model. The 11 metastable basins identified by PCCA with the corresponding representative structures and the transitions probabilities among them are shown. The nodes of the tetrasome (T, T_L_, T_R_), left hexasome (H_L0_, H_L1_, H_L2_), right hexasome (H_R0_, H_R1_, H_R2_, H_R3_), and nucleosome (N) basins, are respectively colored in yellow, blue, cyan and red. The areas of the nodes are proportional to their equilibrium populations. The transition probabilities between basins are represented by the thickness of arrows and the numbers associated (e.g., 3e-2 stands for 3×10^−2^).

### Asymmetric dynamics of 601 nucleosomes

Our MSM also reveals a clear asymmetry between the left and right side of the 601 sequence. For example, the left hexasome state H_L1_ has an equilibrium probability about 20 times higher than its right counterpart H_R1_, consistent to what observed in past experiments on hexasomes [43]. Kinetics itself is also highly asymmetric: binding always occurs with a higher rate on the left side of 601, whereas unbinding occurs with a higher rate on the right (Fig 2), as observed in past experimental studies [17]. For instance, a tetrasome T transitions into a left hexasome H_L1_ after 4×10^6^ MD steps with probability of ∼3×10^−3^, one order of magnitude greater than the probability to observe a T to H_R1_ transition (∼2×10^−4^). Similarly, the rate of nucleosome formation is ten times faster when this last assembly step involves H2A/H2B binding on the left side (i.e., from H_R1_ to N).

What is the molecular origin of this kinetic asymmetry? As shown in Fig 1A, the left side of 601 contains periodically spaced A/T base steps at most 5+10n positions from the dyad, whereas on the right side this periodic pattern is much weaker. The importance of the periodic motifs in nucleosome folding stems from the intrinsic bending of the DNA helix, which lowers the energy cost of wrapping around histones [32,38,44]. Indeed, we find that when the histone octamer is assembled (state N) the left side of the 601 DNA has a much higher probability to be fully wrapped than the right side (Fig 3A). Furthermore, even the distributions of H2A/H2B dimer positions along DNA when they are unbound from the tetramer display strong differences (Fig 3B): the left distribution has two sharp peaks separated by 10 base pairs, one of which corresponds to the location found in the left hexasome and complete nucleosome states, while H2A/H2B positioning on the right side of 601 is much more uniform. This shows that the sequence motifs on the left side of 601 (highlighted again in Fig 3B) facilitate the positioning of the H2A/H2B dimer so that it is poised for a successful binding with the H3/H4 tetramer.

**Fig 3.**
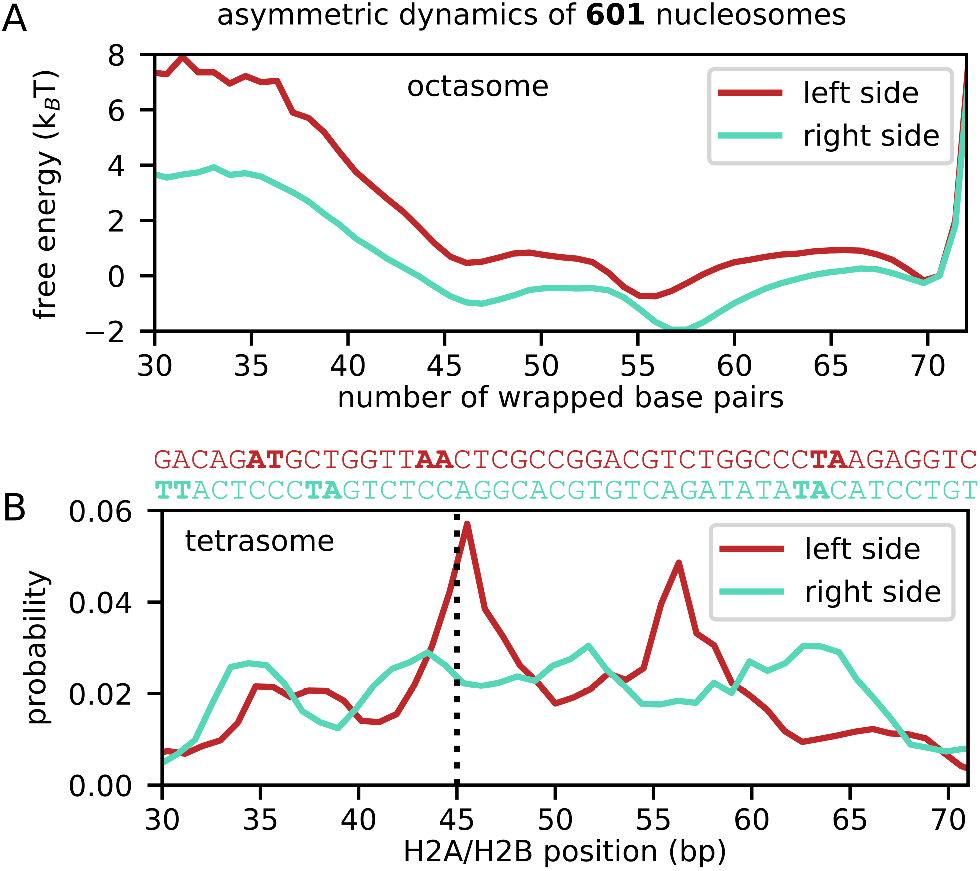
Asymmetric dynamics in 601 nucleosomes at 400 mM. (A) Free energy curves of DNA wrapping for the left (in red) and right (in blue) halves of assembled nucleosome. Only the structural ensemble corresponding to the state N, i.e., complete nucleosome, in the MSM (Fig 2) is used. The free energy at the complete wrapping of 70 bp is set to zero. (B) For the tetrasome states including T, T_R_ and T_L_, we plot the probability of the positions of the unbound H2A/H2B dimers on the left (in red) and the right (in blue) sides of the 601 DNA. The DNA position corresponds to the base pair of the phosphate group that is closest to residue ARG29 on the H2A histone (as described in Methods). On the left side the dimer has a much higher probability to be localized at the same position as when bound to the H3/H4 tetramer (indicated by the black dotted line), facilitating successful assembly. On top of the graph, we align each side of the 601 sequence to the DNA position axis (left side in red, right side in blue).

### Assembly pathways and salt dependence

The analysis of transition pathways [21] (S3 Fig) in the coarse-grained MSM of the 11 long-lived states shows that two pathways make up most of the successful assembly transitions between the tetrasome T and the full nucleosome N: 66% of the pathways go through the more stable left hexasome H_L1_ (T→ H_L1_→ N), while 19% go through the less stable right hexasome H_R1_ (T→ H_R1_→ N). Therefore, all other hexasome conformations (H_L0_, H_L2_, H_R0_, H_R2_ and H_R3_) do not significantly contribute to successful assembly and may be therefore considered off-pathway kinetic traps.

The above landscape of nucleosome assembly and disassembly was obtained at an intermediate ionic strength (400 mM). However, an interesting question is how the assembly and disassembly processes are affected by changes in salt concentration. As described before, our assembly simulations from the tetrasome configurations at 200 mM, 300 mM, and 400 mM salt resulted in a complete nucleosome assembly in 20, 7 and 1 cases out of 215 runs, suggesting that low salt favors assembly. This is reasonable, since nucleosome are stabilized by protein-DNA electrostatic interactions. However, it does not explain the observation that the optimal procedure to assemble nucleosomes *in vitro* is by slowly decreasing the salt concentration from high to low values [23], and otherwise chromatin remodelers are required to successfully complete assembly [14]. One possibility is that at low ionic strength the kinetic landscape becomes more rugged, and the system struggles to escape from off-pathway traps. Indeed, Fig 4A indicates that although lower salt concentration increases the probability to reach the complete nucleosome, it also significantly increases the probability to reach the kinetic traps identified from the MSM (H_L0_, H_L2_, H_R0_, H_R2_ and H_R3_). To further study this effect, we run for each ionic strength 100 10^7^- steps simulations starting from a H_R2_ hexasome conformation, one of the traps that do not participate in the assembly, where the unbound H2A/H2B dimer bridges both left and right sides of DNA. Consistently with our hypothesis, the time required to escape from the initial state increases as we lower the salt concentration (Fig 4B, see Methods for the details of this analysis).

**Fig 4.**
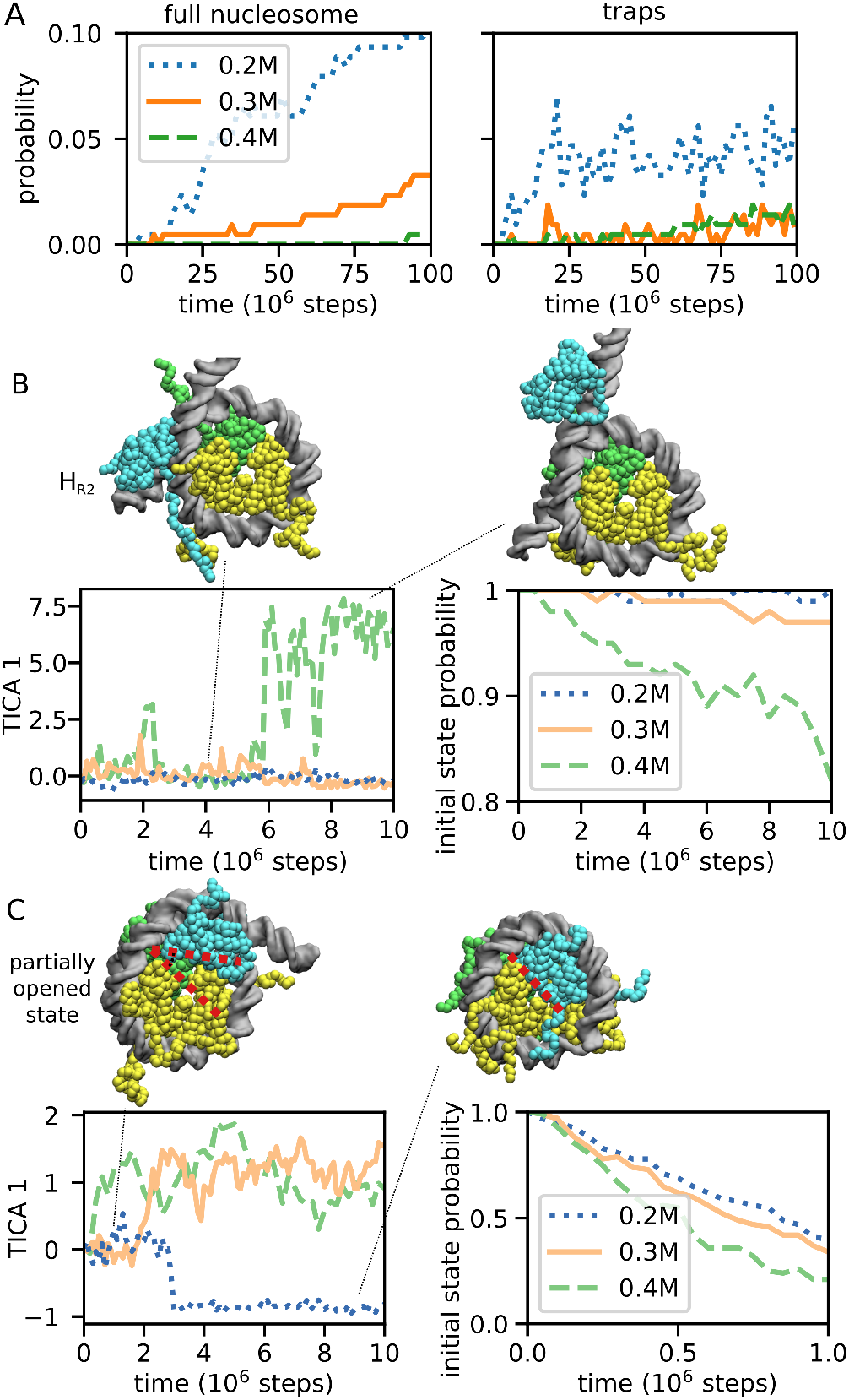
Salt-dependent assembly kinetics and escape from two traps at the salt 200, 300, and 400 mM. (A) The time-dependent probability that, starting from a tetrasome, the system reaches either a full nucleosome (left panel) or one of the kinetic traps (right panel), which are defined as all the off-pathway conformations corresponding to the H_R0_, H_L0_, H_R2_, H_L2_ and H_R3_ basins. (B) The escape kinetics from the H_R2_ kinetic trap. The left panel plots the timeline of the slowest characteristic coordinate (TICA 1, see Supplementary Methods for its definition) for a representative trajectory for each salt concentration. The right panel shows, as a function of time, the probability to remain the initial state, defined as the TICA 1 interval from -4 to +4, obtained from 100 trajectories (see Supplementary Methods for more details). (C) The escape from the partially opened right hexasome identified during the successful hexamer-to-nucleosome assembly events. In order to highlight the conformational change, we indicate with two dotted red lines the wider angle between the unbound H2A/H2B dimer and the neighboring H3/H4 dimer, relative to that found in the fully assembled nucleosome. The left and right panels are the same plot as (B) except that the initial state is defined as the TICA 1 interval from -0.5 to 0.5. The time required to escape from these trapped states increases at low salt.

During the successful assembly trajectories at 200 mM and 300 mM salt, we also found evidence of new metastable states that are not clearly captured by our MSM at 400 mM. In these conformations the nucleosome is already very compact but partially opened: due to the absence of DNA bending at super-helical location +/-3 (i.e., 3 DNA helical turns away from the dyad), one or both H2A/H2B dimers are unbound from the tetramer and oriented at a wider angle relative to the canonical conformation (see snapshot in Fig 4C). During our trajectories, we found this angle to vary between ∼10 and ∼50 degrees (the angle was computed based on the orientation of the longest helices in histones H2A and H4). PDB coordinates of representative partially opened conformations (part of the H_L_, H_R_ or T states) have been deposited in S1 Data together with the other metastable states defined from the MSM. To further investigate their properties, we run 100 10^7^-steps simulations starting from the partially opened right hexasome at 200, 300, and 400 mM salt. Fig 4C shows that, similarly to the H_R2_ state, the escape from the partially opened state becomes slower as we decrease the salt concentration, presumably because this conformation is stabilized by electrostatic interactions of the H2A C-terminal and H2B N-terminal histone tails bridging the two DNA gyres. Furthermore, at 300 mM we find that starting from a partially opened hexasome conformation the probability to transition either into the complete nucleosome or into a more open hexasome is about 50%, similarly to a transition state (within 5×10^6^ MD steps, we observe complete assembly events in 269 runs out of 400 starting from the left partially opened state, and in 156 out of 400 from the right one). Finally, we note that all the evidence points to these partially opened nucleosomes being the same as those identified in a recent FRET study [17]. In both experiments and in our simulations, 1. the angle between the open H2A/H2B dimer and the tetramer very similar (a value of 20 degrees was suggested from experiments), 2. the stability of these states decreases with increasing ionic strength, and 3. they are intermediates along the nucleosome assembly pathways.

In order to explore whether the disassembly mechanism is also affected by the salt concentration, we run additional 10^8^-step simulations starting from the complete nucleosome at 500, 600, 700, and 800 mM salt (30 trajectories each). As expected, nucleosomes become unstable at high salt, with frequent DNA unwrapping and dimer unbinding. Interestingly, disassembly pathways appear qualitatively different at intermediate (400 mM) and high (800 mM) salt: while in the former case H2A/H2B unbinding from the tetramer proceeds cooperatively with the unwrapping of DNA, in the latter case DNA unwrapping anticipates histone unbinding (Fig 5). This trend is in qualitative agreement with the experimental literature: the cooperative transition is consistent with the recent results obtained by FRET at intermediate salt [17], while the non-cooperative behavior was suggested by a combination of SAXS and FRET at very high 1.9 M salt concentration [9].

**Fig 5.**
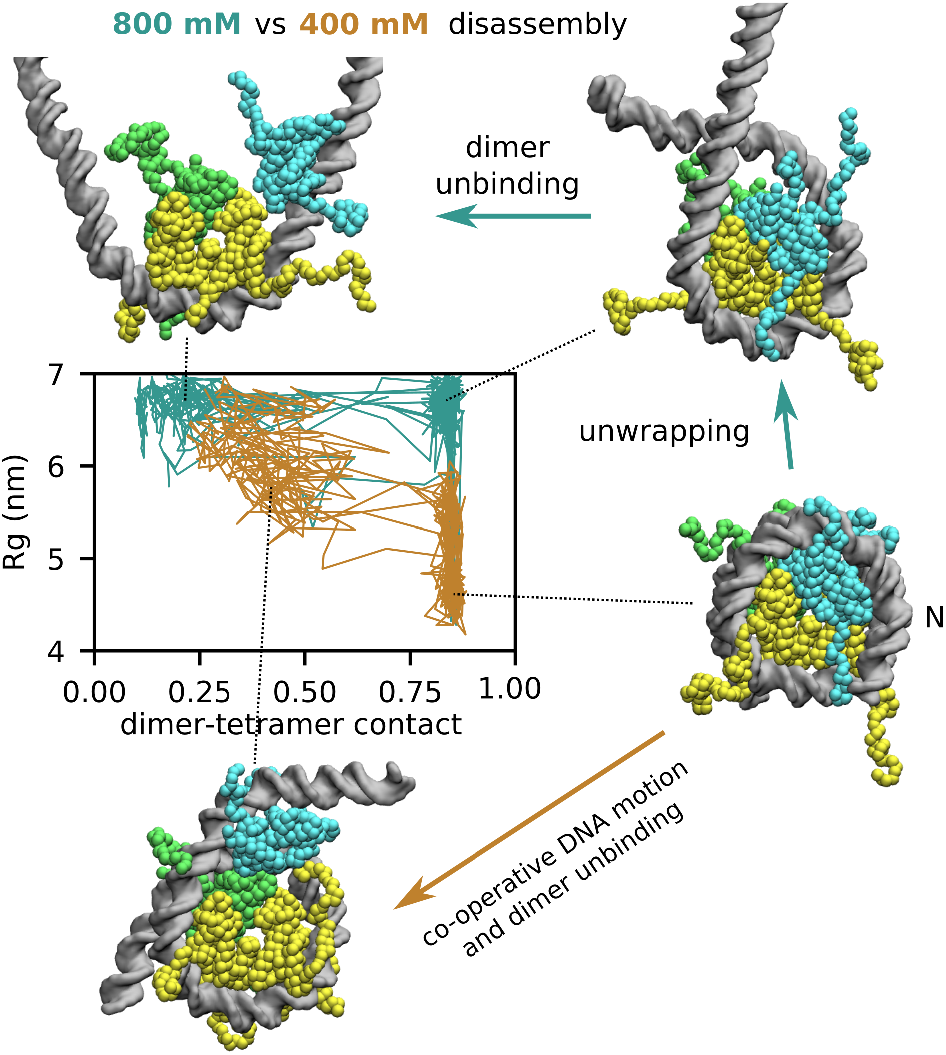
Salt-dependent nucleosome disassembly pathways. We project a set of representative disassembly trajectories along the DNA radius of gyration (Rg, the vertical axis) and a coordinate capturing dimer-tetramer contact formation (the horizontal axis, see Supplementary Methods for the precise definition), at either 400 mM (in brown) or 800 mM salt (in cyan). While at intermediate salt disassembly occurs in a single step involving dimer unbinding from the tetramer cooperatively with the stretching motion of DNA, at high salt dimer unbinding follows a significant unwrapping of DNA from the histone octamer.

## Discussion

In conclusion, our coarse-grained MD simulations revealed the complex landscape of nucleosome assembly and its modulation by salt and DNA sequence. Firstly, we used Markov state modeling to summarize the assembly dynamics on 601 sequences at 400 mM salt concentration (at which the assembly is a reversible). Our MSM highlighted, apart from the expected canonical nucleosome, tetrasome and hexasome states, several long-lived configurations characterized by alternative interactions between histones and DNA. Interestingly, in one of the right hexasome states (H_R3_), a canonical histone hexamer wraps the DNA in a right-handed way. Right-handed nucleosomes can form in experiments after the application of positive DNA supercoiling [45,46] and are also found *in vivo* within centromeres [15]. However, while past works investigated how such change in DNA handedness could originate from an equivalent change in the arrangement of the histone folds [45–47], our simulations suggest that these right-handed structures may also form via an alternative wrapping of the DNA around a canonical histone hexamer, without requiring a complex rearrangement of histones [45]. Since this state is found only on one side of 601, the simulations further suggest that right-handed nucleosomes are stabilized by specific DNA sequences. Torsional stresses, such as those generated by the passage of RNA polymerase, are also expected to affect the stability of various nucleosome configurations, as found in experiments on nucleosomes with super-coiled DNA [48], and in a recent MD study that investigated the effect of torsion on nucleosomal DNA unwrapping [49].

Our simulations also provide fresh insights into asymmetric nucleosome dynamics on 601 DNA. This sequence is characterized by several nucleosome positioning motifs, namely A/T base steps periodically spaced every 10 bp, which favor the wrapping of DNA around histones. However, these motifs are asymmetrically distributed along the sequence, favoring H2A/H2B dimer assembly on the left side of 601 by about one order of magnitude. Asymmetric disassembly of nucleosomes along 601 was observed in many experiments [9,17,38]; our MD simulations revealed that DNA sequence motifs promote assembly by directing the binding of H2A/H2B dimers towards the H3/H4 target, an effect originating from the co-operativity between histone binding and DNA bending at low salt concentration. We speculate that by promoting the assembly of nucleosomes directly at highly flexible genomic regions [50], while avoiding stiff DNA regions such as poly-A tracts, the sequence dependence in the rates of assembly may also favor the maintenance of nucleosome positioning *in vivo* after disruptive events such as replication. In the future, we plan to investigate the dynamics of nucleosomes on genomic sequences to further explore this hypothesis.

Finally, our simulations highlighted how salt dependence can have a large effect on the kinetics of the system. The MSM generated at 400 mM salt shows that successful nucleosome assembly pathways are relatively simple: the dominant pathway from the tetrasome (T) to the complete nucleosome conformation (N) goes through a single intermediate hexasome state with a H2A/H2B dimer bound to the H3/H4 tetramer on the left side of 601 (H_L1_), while the second dominant pathway goes through the opposite intermediate hexasome (H_R1_) where the dimer is bound to the tetramer on the right side of 601. All other metastable states do not significantly participate into the assembly process, effectively acting as kinetic traps slowing down nucleosome formation. While low salt favors assembly, we find that it also stabilizes off-pathway kinetic traps, explaining the importance of salt dialysis for the successful assembly process [23]. Under some conditions, experiments found that histones and DNA form non-canonical pre-nucleosome structures that require chromatin remodelers to be converted into the canonical forms [14]. We speculate that pre-nucleosomes may resemble the haxasome states H_L2_ and H_R2_, where the unbound H2A/H2B dimer bridges the two opposite ends of nucleosomal DNA to prevent successful assembly unless some large rearrangement first occurs.

Simulations at 200 and 300 mM also revealed critical intermediate states along the successful pathways of nucleosome assembly: here the nucleosome is almost completely formed, but one H2A/H2B dimer does not yet make direct contacts with the H3/H4 tetramer, with the two interfaces at a wider angle relative to the canonical nucleosome. These partially opened nucleosomes where also recently identified in a series of FRET experiments [17]. Simulations indicate that these partially opened states are roughly halfway along the nucleosome pathways at low salt (i.e., they are transition states); in virtue of this, modulating the stability of this state, for instance by histone chaperones or epigenetic modifications, should have a large effect on assembly and disassembly rates, suggesting its potential role *in vivo*. Again, we found that the formation of these structures does not require any conformational change within histones (which may nevertheless play a role in nucleosome dynamics [51]), but are instead stabilized by non-specific electrostatic interactions between histone tails and DNA. In addition to conformational stability, salt concentration also affects the assembly/disassembly pathway itself: while at low and intermediate salt the H2A/H2B dimer binds the H3/H4 tetramer co-operatively with DNA bending (so that DNA sequence can directly affect kinetics), at high salt these two processes are uncoupled, with DNA unwrapping anticipating dimer-tetramer dissociation. These results recapitulate past experimental observations under various conditions [9,17].

The position coordinates of the metastable nucleosome conformations deposited in the supplementary material will represent a useful resource for researchers interested in further investigating nucleosome assembly, for instance by aiding the design of FRET experiments [17], or by providing starting configurations for all-atom MD simulations through a back-mapping procedure [42]. Finally, we envision that our coarse-grained model will be of great use to further characterize nucleosome assembly in more realistic scenarios typical of *in vivo* experiments, for example in the presence of RNA polymerase [52], histone chaperones [47], and remodelers [13].

## Methods

### Coarse-grained model

To enable the complete study of nucleosome assembly and disassembly, we utilize coarse-grained molecular modeling: we map each amino-acid to a single bead [25], and each nucleotide to three beads corresponding to base, sugar, and phosphate groups [24]. The functional potentials and force-field parameters are chosen according to the AICG2+ structure-based model for histones [25] and the 3SPN2.C sequence-dependent model for DNA [24]. The flexible histone tails (residue ids 1-32 for H3, 1-23 for H4, 1-14 and 121-128 for H2A, 1-26 for H2B) are modeled according to a statistical local specifically designed for disordered regions [53]. The potential of the remaining folded regions is based on the structures of the first copies of each H3/H4 and H2A/H2B dimers present in the nuclesome crystal structure with PDB id 1KX5 (using a single reference for each pair of dimers ensures the symmetry of the histone octamer, which is important to study the asymmetry induced by DNA sequence). Briefly, whenever two protein residues are within a certain cutoff distance in the crystal structure, the coarse-grained potential includes an attractive interaction that favors the binding of the two residues [25].

Experimentally, in the absence of DNA or at high salt the histone octamer breaks into a H3/H4 tetramer and two H2A/H2B dimers [9,17,54], but using the default AICG2+ potential we do not observe histone octamer breakage under these conditions. This happens because the default AICG2+ scaling factor defining the overall strength of the residue-residue interactions was optimized for studying the folding of single-domain proteins, but in our case the attraction between the H3/H4 and H2A/H2B dimers is over-estimated. In order to find a suitable scaling factor for the native interactions between H3/H4 and H2A/H2B residues, we scanned different values performing MD simulation at either low salt (200 mM), at which we expect our nucleosomes to be stable, or high salt (800 mM), at which we expect disassembly. We find that scaling factors lower than 0.2 lead to unstable nucleosomes even at low salt, whereas for values higher than 0.4 we do not observe disassembly at even at high salt. Rescaling the dimer-tetramer interactions by a factor of 0.3 reproduces the expected behavior of nucleosomes: out 30 independent simulation runs of 10^8^ MD steps starting from a fully assembled nucleosome, we observed no disassembly events at 200 mM salt, whereas we observed disassembly in 4 out of 30 runs at 800 mM; as shown by the MSM reported in the Results section, nucleosome assembly is reversible at intermediate 400 mM salt, with comparable equilibrium populations of tetrasomes and complete nucleosomes. We use the dimer-tetramer scaling factor of 0.3 in all simulations reported in this study.

Histones and DNA interact via excluded volume, Debye–Hückel electrostatics, and hydrogen bond interactions. The hydrogen bonds are modeled according to a distance- and angle-dependent potential based on nucleosome crystal structures, and was calibrated based on nucleosome structural stability and DNA unwrapping [26]. The size of the beads to define excluded volume interactions, and the parameter settings for the hydrogen bonds are the same used in our previous works [26–28]. The charges on the histone residues are determined via the RESPAC method [55], which optimizes the coarse-grained electrostatic potential against the all-atom one. The RESPAC procedure was applied to the globular part of the H3/H4 tetramer and the two H2A/H2B dimers, whereas for the flexible tails we used the standard integer charges on the Lysine, Arginine, Aspartic and Glutamic acids residues only.

### Simulation setup

All reported molecular dynamics (MD) simulations have been performed with the software Cafemol [56] by integrating the equations of motion using Langevin dynamics and default MD settings. During the MD simulations, we apply harmonic restraints on the structured region of the two H3 histones (0.001 kcal/mol/Å^2^) and on the phosphate beads of the DNA dyad residues (bp id 73, 0.1 kcal/mol/Å^2^), limiting large-scale sliding of DNA relative to the H3/H4 tetramer (but not the unbound H2A/H2B dimers). The nucleosome sliding mechanism was already analyzed in previous studies [26,27], and especially for the 601 positioning sequence considered here can be extremely slow. In our simulations, the H3/H4 is always placed at its optimal experimental position on the DNA sequence; preventing its sliding allows to avoid exploring less favorable positions and to focus solely on the assembly dynamics, which is our main focus. Finally, the system is enclosed into a spherical volume with a radius of 20 nm by a harmonic potential. Cafemol input files to run nucleosome assembly simulations are provided in the supplementary S2 Data.

The starting conformations for the 1000 MD simulation runs at 400 mM used to generate the MSM have been obtained during high-salt (800 mM) simulations starting from complete nucleosomes, and then selecting 1000 timeframes that include tetrasomes (215 conformations), left and right hexasomes (105 and 118 conformations, respectively), and nucleosomes (562 conformations). To quantify the salt-dependence of the nucleosome kinetics, we also performed, at both 200 and 300 mM salt, 215 10^8^-steps simulation runs starting from the 215 tetrasome conformations. To analyze the escape from 2 metastable states (either the H_R2_ hexasome or the partially opened state) as a function of salt concentration, we run 100 independent 10^7^-steps trajectories starting from the same conformation (either H_R2_ or partially opened H_R1_) at 200, 300, and 400 mM salt. In order to characterize the time necessary to escape from these states (analysis in Fig 4), we projected the molecular positions onto the slowest TICA motions [41] within each basin. The TICA projection was performed separately for the H_R2_ and partially opened metastable basins based on the positions of H2A residues 46 to 73 of the unbound dimer and the left side of DNA (the one interacting with the unbound H2A/H2B dimer) during the simulations at 300 mM salt.

### Markov state modeling and analysis of metastable states

To generate the Markov state model of nucleosome assembly at 400 mM salt from our 1000 MD simulations, we first have to cluster the configurations into discrete states based on a distance between configurations defined on a suitable space. To this aim, we project the particle positions onto a series of collective variables that capture all the relevant modes of nucleosome dynamics: H2A/H2B dimer binding to DNA and to H3/H4, DNA wrapping, and DNA handedness. Since the two H2A/H2B dimers are identical, we generated variables that are symmetric under an exchange of these two dimers. In principle, the two H3/H4 binding interfaces are also identical, but the 601 DNA that is wrapped around the tetramer is not symmetric, so we do want to distinguish between the left and right H3/H4 interfaces, corresponding respectively to the dimer binding the left or the right side of the 601 DNA. We defined 5 types of collective variables (for a total of 4+4+2+2+1=13 collective variables):

- For each H3/H4 tetramer interface, the sum and the maximum value of the contacts with the two H2A/H2B dimers. A dimer-tetramer contact is defined as c=1/(1+d/σ), where σ=1 nm and d is the distance between the center of mass of H2B residues 73-100 and the center of mass of H4 residues 65-92 (corresponding to the two interfacial regions). Therefore, a contact takes a value close to 1 when a H2A/H2B dimer binds the H3/H4 tetramer in its native configuration, whereas it is close to 0 when the H2A/H2B dimer is far from the H3/H4 tetramer. We first compute the 4 contacts between each of the 2 tetramer interfaces (t1 and t2) with the two dimers (d1 and d2): c_t1,d1_, c_t1,d2_, c_t2,d1_, and c_t2,d2_. Then, we define the 4 collective variables: c_t1,d1_+c_t1,d2_; c_t2,d1_+c_t2,d2_; max(c_t1,d1_,c_t1,d2_); and max(c_t2,d1_,c_t2,d2_).
- For each side of the DNA (left, residue ids 1-73, and right, ids 75-147), the sum and the maximum value of the contacts with the two H2A/H2B dimers. A DNA-dimer contact is defined as 1/(1+d/σ), where σ=1nm, and d is the distance between residue ARG29 on H2A (in the canonical nucleosome this residue binds the DNA at SHL +/- 4.5) and the closest DNA phosphate group. The sum of the DNA-dimer contacts enables us to distinguish between the various tetrasome conformations where two dimers can bind either the same (states T_L_ and T_R_ in Fig 2 of the main text) or two different DNA regions (state T). The maximum value of the contact distinguishes strong or weak binding of the H2A/H2B dimer to the DNA.
- For each side of the DNA (left and right), the H2A/H2B dimer location along the DNA, which is defined as the residue id of the DNA phosphate group (on the 5’ strand for either side) that is closest to the H2A ARG29 residue on the dimer closest to the considered DNA side. For example, for the location on the left side of the DNA, if the first H2A/H2B dimer is closer than the second one (based on the minimum distance above), we consider the base pair id of the phosphate group closest to the first dimer. This variable enables us to capture the sliding of the H2A/H2B dimers along the DNA when they are unbound from the H3/H4 tetramer.
- For each side of the DNA, the number of base pairs wrapped around the histones, where a base pair is considered unwrapped if the center of mass of its base groups is displaced radially from the nucleosome super-helix axis by more than 6 Å.
- The handedness of the nucleosome, which is captured by the component of the distance vector between the centers of mass of base pairs with ids -35 and +35 (relative to the dyad id 0) along the nucleosome super-helix axis. Positive values correspond to canonical left-handed nucleosomes, whereas negative ones correspond to right-handed conformations such as those in the metastable state H_R3_ in Fig 2 of the main text.

Starting from these 13 collective variables, we aggregate all 1000 MD trajectories and perform a further dimensionality reduction using time-lagged independent component analysis (TICA) [41] as implemented in PyEMMA 2 [57] using a lag-time of 10^6^ MD steps. We then select the top 8 slowest coordinates (which explain 95% of the kinetic variance in the data) and cluster the conformations into 400 discrete states using k-means [58]. The MSM was then generated by PyEMMA 2 [57] using a lag-time of 4×10^6^ MD steps. Further increasing the lag-time does not lead to significant changes in the implied time scales of the MSM, indicating the robustness of the results (S1 Fig). The TICA co-ordinates are also ideally suited for visualization purposes: in S2 Fig we show the first two coordinates are sufficient to separate the system into 4 regions corresponding to nucleosomes, left and right hexasomes and tetrasomes.

## Supporting Information Captions

S1 Fig. Implied time scales of the MSM as a function of the chosen lag-time. For the final model, we use a lag-time of 4×10^6^ MD steps, after which most of the implied time scales have essentially converged. The shaded regions indicate 95 confidence intervals.

S2 Fig. Nucleosome conformations projected along the first two slowest TICA coordinates and colored according to the PCCA metastable basin (see main text). These coordinates already separate the system into 4 distinct regions corresponding to complete nucleosomes, left and right hexasomes, and tetrasomes. The first TICA is proportional to the number of contacts between the H3/H4 tetramer and the H2A/H2B dimers, whereas the second TICA is proportional to the difference between the right and left dimer-tetramer contacts. More TICA coordinates are necessary to further separate these 4 regions into all the metastable basins.

S3 Fig. Analysis of the transition pathways. We show the metastable basins and the two top pathways to reach the complete nucleosome N starting from the tetrasome T. Both pathways involve a single intermediate state corresponding to one of the two caninical hexasomes (H_L1_ or H_R1_). Above the arrows we indicate the percentage of pathways passing through this transition. The x coordinate corresponds to the committor probability as computed from the MSM, which is defined as the probability to reach state N before coming back to state T.

S1 Movie. Representative assembly trajectory at 400 mM starting from a tetrasome conformation, the same shown in Fig 1B in the main text.

S1 Data: For each of the 11 metastable states identified from our MSM at 400 mM using PCCA clustering, we deposited 4 representative coarse-grained nucleosome conformations in PDB format. The conformations are named “state*conf#.pdb”, where * corresponds to the state name as described in the text (T, T_L_, T_R_, H_L0_, H_L1_, H_L2_, H_R0_, H_R1_, H_R2_ and H_R3_, N), and # goes from 1 to 4. The data also contain 4 representative partially opened conformations for each of the left hexasome, right hexasome, and tetrasome states, named “partially opened_*_conf#.pdb”, where * is H_L_, H_R_ or T and # goes from 1 to 4.

S2 Data: Example Cafemol input files used to run nucleosome assembly simulations as described in the Methods section.

## References

1. Luger K, Mäder AW, Richmond RK, Sargent DF, Richmond TJ. Crystal structure of the nucleosome core particle at 2.8 Å resolution. Nature. 1997;389: 251–260. doi:10.1038/38444

2. Zhou K, Gaullier G, Luger K. Nucleosome structure and dynamics are coming of age. Nat Struct Mol Biol. 2018; 1. doi:10.1038/s41594-018-0166-x

3. Polach KJ, Widom J. Mechanism of Protein Access to Specific DNA Sequences in Chromatin: A Dynamic Equilibrium Model for Gene Regulation. J Mol Biol. 1995;254: 130–149. doi:10.1006/jmbi.1995.0606

4. Small EC, Xi L, Wang J-P, Widom J, Licht JD. Single-cell nucleosome mapping reveals the molecular basis of gene expression heterogeneity. Proc Natl Acad Sci U S A. 2014;111: E2462–71. doi:10.1073/pnas.1400517111

5. Venkatesh S, Workman JL. Histone exchange, chromatin structure and the regulation of transcription. Nat Rev Mol Cell Biol. 2015;16: 178–189. doi:10.1038/nrm3941

6. Michieletto D, Chiang M, Colì D, Papantonis A, Orlandini E, Cook PR, et al. Shaping epigenetic memory via genomic bookmarking. Nucleic Acids Res. 2018;46: 83–93. doi:10.1093/nar/gkx1200

7. Kireeva ML, Walter W, Tchernajenko V, Bondarenko V, Kashlev M, Studitsky VM. Nucleosome Remodeling Induced by RNA Polymerase II: Loss of the H2A/H2B Dimer during Transcription. Mol Cell. 2002;9: 541–552. doi:10.1016/S1097-2765(02)00472-0

8. Serra-Cardona A, Zhang Z. Replication-Coupled Nucleosome Assembly in the Passage of Epigenetic Information and Cell Identity. Trends Biochem Sci. 2017;43: 136–148. doi:10.1016/J.TIBS.2017.12.003

9. Chen Y, Tokuda JM, Topping T, Meisburger SP, Pabit SA, Gloss LM, et al. Asymmetric unwrapping of nucleosomal DNA propagates asymmetric opening and dissociation of the histone core. Proc Natl Acad Sci U S A. 2017;114: 334–339. doi:10.1073/pnas.1611118114

10. Wilhelm FX, Wilhelm ML, Erard M, Daune MP. Reconstitution of chromatin: assembly of the nucleosome. Nucleic Acids Res. 1978;5: 505–521. doi:10.1093/nar/5.2.505

11. Fathizadeh A, Berdy Besya A, Reza Ejtehadi M, Schiessel H. Rigid-body molecular dynamics of DNA inside a nucleosome. Eur Phys J E. 2013;36: 21. doi:10.1140/epje/i2013-13021-4

12. Andrews AJ, Chen X, Zevin A, Stargell LA, Luger K. The Histone Chaperone Nap1 Promotes Nucleosome Assembly by Eliminating Nonnucleosomal Histone DNA Interactions. Mol Cell. 2010;37: 834–842. doi:10.1016/J.MOLCEL.2010.01.037

13. Clapier CR, Iwasa J, Cairns BR, Peterson CL. Mechanisms of action and regulation of ATP-dependent chromatin-remodelling complexes. Nat Rev Mol Cell Biol. 2017;18: 407–422. doi:10.1038/nrm.2017.26

14. Fei J, Torigoe SE, Brown CR, Khuong MT, Kassavetis GA, Boeger H, et al. The prenucleosome, a stable conformational isomer of the nucleosome. Genes Dev. 2015;29: 2563–2575. doi:10.1101/gad.272633.115

15. Furuyama T, Henikoff S. Centromeric nucleosomes induce positive DNA supercoils. Cell. 2009;138: 104–113. doi:10.1016/j.cell.2009.04.049

16. Kubik S, Bruzzone MJ, Jacquet P, Falcone J-L, Rougemont J, Shore D. Nucleosome Stability Distinguishes Two Different Promoter Types at All Protein-Coding Genes in Yeast. Mol Cell. 2015;60: 422–434. doi:10.1016/J.MOLCEL.2015.10.002

17. Gansen A, Felekyan S, Kühnemuth R, Lehmann K, Tóth K, Seidel CAM, et al. High precision FRET studies reveal reversible transitions in nucleosomes between microseconds and minutes. Nat Commun. 2018;9: 4628. doi:10.1038/s41467-018-06758-1

18. Rhee HS, Bataille AR, Zhang L, Pugh BF. Subnucleosomal structures and nucleosome asymmetry across a genome. Cell. 2014;159: 1377–1388. doi:10.1016/j.cell.2014.10.054

19. Brehove M, Shatoff E, Donovan BT, Jipa CM, Bundschuh R, Poirier MG. DNA sequence influences hexasome orientation to regulate DNA accessibility. Nucleic Acids Res. 2019;47: 5617–5633. doi:10.1093/nar/gkz283

20. Lowary P., Widom J. New DNA sequence rules for high affinity binding to histone octamer and sequence-directed nucleosome positioning. J Mol Biol. 1998;276: 19–42. doi:10.1006/jmbi.1997.1494

21. Noé F, Schütte C, Vanden-Eijnden E, Reich L, Weikl TR. Constructing the equilibrium ensemble of folding pathways from short off-equilibrium simulations. Proc Natl Acad Sci U S A. 2009;106: 19011–19016. doi:10.1073/pnas.0905466106

22. Basu A, Bobrovnikov DG, Qureshi Z, Kayikcioglu T, Ngo TTM, Ranjan A, et al. Measuring DNA mechanics on the genome scale. Nature. 2021;589: 462–467. doi:10.1038/s41586-020-03052-3

23. Lee K-M, Narlikar G. Assembly of Nucleosomal Templates by Salt Dialysis. Current Protocols in Molecular Biology. Hoboken, NJ, USA: John Wiley & Sons, Inc.; 2001. pp. 21.6.1-21.6.16. doi:10.1002/0471142727.mb2106s54

24. Freeman GS, Hinckley DM, Lequieu JP, Whitmer JK, de Pablo JJ. Coarse-grained modeling of DNA curvature. J Chem Phys. 2014;141: 165103. doi:10.1063/1.4897649

25. Li W, Wang W, Takada S. Energy landscape views for interplays among folding, binding, and allostery of calmodulin domains. Proc Natl Acad Sci U S A. 2014;111: 10550–10555. doi:10.1073/pnas.1402768111

26. Niina T, Brandani GB, Tan C, Takada S. Sequence-dependent nucleosome sliding in rotation-coupled and uncoupled modes revealed by molecular simulations. PLOS Comput Biol. 2017;13: e1005880. doi:10.1371/journal.pcbi.1005880

27. Brandani GB, Niina T, Tan C, Takada S. DNA sliding in nucleosomes via twist defect propagation revealed by molecular simulations. Nucleic Acids Res. 2018;46: 2788– 2801. doi:10.1093/nar/gky158

28. Brandani GB, Takada S. Chromatin remodelers couple inchworm motion with twist-defect formation to slide nucleosomal DNA. PLOS Comput Biol. 2018;14: e1006512. doi:10.1371/journal.pcbi.1006512

29. Lequieu J, Schwartz DC, de Pablo JJ. In silico evidence for sequence-dependent nucleosome sliding. Proc Natl Acad Sci U S A. 2017;44: E9197–E9205. doi:10.1073/pnas.1705685114

30. Kenzaki H, Takada S. Partial Unwrapping and Histone Tail Dynamics in Nucleosome Revealed by Coarse-Grained Molecular Simulations. PLOS Comput Biol. 2015;11: e1004443. doi:10.1371/journal.pcbi.1004443

31. Lequieu J, Córdoba A, Schwartz DC, de Pablo JJ. Tension-Dependent Free Energies of Nucleosome Unwrapping. ACS Cent Sci. 2016;2: 660–666. doi:10.1021/acscentsci.6b00201

32. Freeman GS, Lequieu JP, Hinckley DM, Whitmer JK, de Pablo JJ. DNA Shape Dominates Sequence Affinity in Nucleosome Formation. Phys Rev Lett. 2014;113: 168101. doi:10.1103/PhysRevLett.113.168101

33. Zhang B, Zheng W, Papoian GA, Wolynes PG. Exploring the Free Energy Landscape of Nucleosomes. J Am Chem Soc. 2016;138: 8126–8133. doi:10.1021/jacs.6b02893

34. Winogradoff D, Aksimentiev A. Molecular Mechanism of Spontaneous Nucleosome Unraveling. J Mol Biol. 2018;431: 323–335. doi:10.1016/J.JMB.2018.11.013

35. Shaytan AK, Armeev GA, Goncearenco A, Zhurkin VB, Landsman D, Panchenko AR. Coupling between Histone Conformations and DNA Geometry in Nucleosomes on a Microsecond Timescale: Atomistic Insights into Nucleosome Functions. J Mol Biol. 2016;428: 221–237. doi:10.1016/j.jmb.2015.12.004

36. Kono H, Sakuraba S, Ishida H. Free energy profiles for unwrapping the outer superhelical turn of nucleosomal DNA. PLOS Comput Biol. 2018;14: e1006024. doi:10.1371/journal.pcbi.1006024

37. Rychkov GN, Ilatovskiy A V., Nazarov IB, Shvetsov A V., Lebedev D V., Konev AY, et al. Partially Assembled Nucleosome Structures at Atomic Detail. Biophys J. 2017;112: 460–472. doi:10.1016/j.bpj.2016.10.041

38. Ngo TTM, Zhang Q, Zhou R, Yodh JG, Ha T. Asymmetric Unwrapping of Nucleosomes under Tension Directed by DNA Local Flexibility. Cell. 2015;160: 1135– 1144. doi:10.1016/j.cell.2015.02.001

39. Deuflhard P, Weber M. Robust Perron cluster analysis in conformation dynamics. Linear Algebra Appl. 2005;398: 161–184. doi:10.1016/j.laa.2004.10.026

40. Noé F, Wu H, Prinz J-H, Plattner N. Projected and hidden Markov models for calculating kinetics and metastable states of complex molecules. J Chem Phys. 2013;139: 184114. doi:10.1063/1.4828816

41. Naritomi Y, Fuchigami S. Slow dynamics in protein fluctuations revealed by time-structure based independent component analysis: The case of domain motions. J Chem Phys. 2011;134: 065101. doi:10.1063/1.3554380

42. Shimizu M, Takada S. Reconstruction of Atomistic Structures from Coarse-Grained Models for Protein-DNA Complexes. J Chem Theory Comput. 2018;14: 1682–1694. doi:10.1021/acs.jctc.7b00954

43. Levendosky RF, Sabantsev A, Deindl S, Bowman GD. The Chd1 chromatin remodeler shifts hexasomes unidirectionally. Elife. 2016;5: e21356. doi:10.7554/eLife.21356

44. Culkin J, de Bruin L, Tompitak M, Phillips R, Schiessel H. The role of DNA sequence in nucleosome breathing. Eur Phys J E. 2017;40: 106. doi:10.1140/epje/i2017-11596-2

45. Lavelle C, Recouvreux P, Wong H, Bancaud A, Viovy J-L, Prunell A, et al. Right-handed nucleosome: myth or reality? Cell. 2009;139: 1216–1217. doi:10.1016/j.cell.2009.12.014

46. Hamiche A, Carot V, Alilat M, De Lucia F, O’Donohue MF, Revet B, et al. Interaction of the histone (H3-H4)2 tetramer of the nucleosome with positively supercoiled DNA minicircles: Potential flipping of the protein from a left-to a right-handed superhelical form. Proc Natl Acad Sci U S A. 1996;93: 7588–7593. doi:10.1073/PNAS.93.15.7588

47. Zhao H, Winogradoff D, Dalal Y, Papoian GA. The Oligomerization Landscape of Histones. Biophys J. 2019;116: 1845–1855. doi:10.1016/j.bpj.2019.03.021

48. Clark DJ, Felsenfeld G. Formation of nucleosomes on positively supercoiled DNA. EMBO J. 1991;10: 387–395. doi:10.1002/j.1460-2075.1991.tb07960.x

49. Ishida H, Kono H. Torsional stress can regulate the unwrapping of two outer half superhelical turns of nucleosomal DNA. Proc Natl Acad Sci. 2021;118: e2020452118. doi:10.1073/pnas.2020452118

50. Segal E, Widom J. Poly(dA:dT) tracts: major determinants of nucleosome organization. Curr Opin Struct Biol. 2009;19: 65–71. doi:10.1016/j.sbi.2009.01.004

51. Bilokapic S, Strauss M, Halic M. Histone octamer rearranges to adapt to DNA unwrapping. Nat Struct Mol Biol. 2018;25: 101–108. doi:10.1038/s41594-017-0005-5

52. Kujirai T, Ehara H, Fujino Y, Shirouzu M, Sekine S, Kurumizaka H. Structural basis of the nucleosome transition during RNA polymerase II passage. Science. 2018;362: 595–598. doi:10.1126/science.aau9904

53. Terakawa T, Takada S. Multiscale ensemble modeling of intrinsically disordered proteins: p53 N-terminal domain. Biophys J. 2011;101: 1450–1458. doi:10.1016/j.bpj.2011.08.003

54. Eickbush TH, Moudrianakis EN. The histone core complex: an octamer assembled by two sets of protein-protein interactions. Biochemistry. 1978;17. Available: URLhttps://pubs.acs.org/doi/pdf/10.1021/bi00616a016

55. Terakawa T, Takada S. RESPAC: Method to Determine Partial Charges in Coarse-Grained Protein Model and Its Application to DNA-Binding Proteins. J Chem Theory Comput. 2014;10: 711–721. doi:10.1021/ct4007162

56. Kenzaki H, Koga N, Hori N, Kanada R, Li W, Okazaki K, et al. CafeMol: A Coarse-Grained Biomolecular Simulator for Simulating Proteins at Work. J Chem Theory Comput. 2011;7: 1979–1989. doi:10.1021/ct2001045

57. Scherer MK, Trendelkamp-Schroer B, Paul F, Pérez-Hernández G, Hoffmann M, Plattner N, et al. PyEMMA 2: A Software Package for Estimation, Validation, and Analysis of Markov Models. J Chem Theory Comput. 2015;11: 5525–5542. doi:10.1021/acs.jctc.5b00743

58. Lloyd S. Least squares quantization in PCM. IEEE Trans Inf Theory. 1982;28: 129– 137. doi:10.1109/TIT.1982.1056489

