## Supplementary figures and images for "The Kinetic Landscape of Nucleosome Assembly: A Coarse-Grained Molecular Dynamics Study"

### S1 Fig

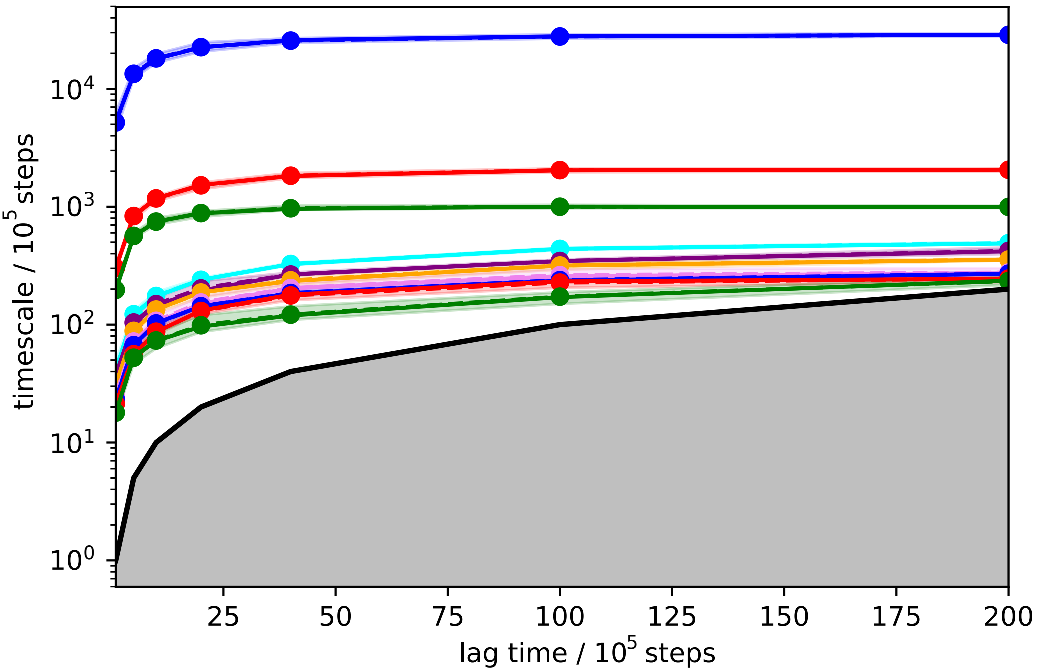

### S2 Fig

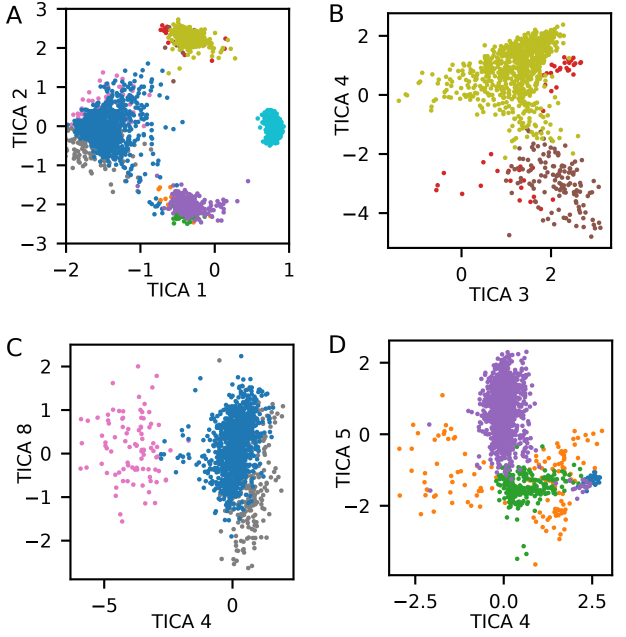

### S3 Fig

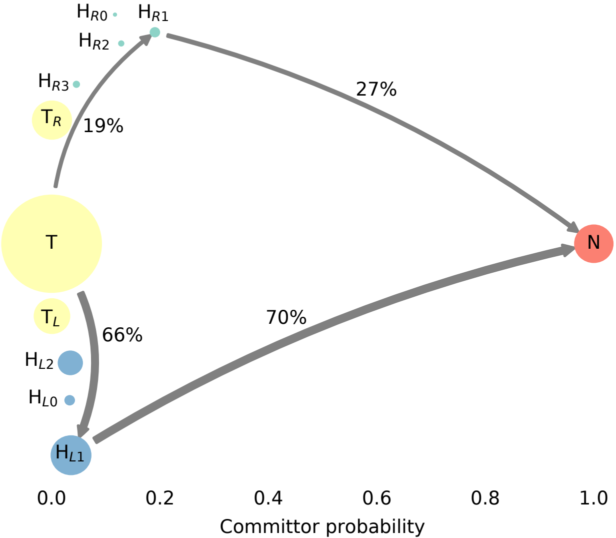
